# An adaptive filtering framework for non-specific and inefficient reactions in multiplex digital PCR based on sigmoidal trends

**DOI:** 10.1101/2022.04.11.487847

**Authors:** Luca Miglietta, Ke Xu, Priya Chhaya, Louis Kreitmann, Kerri Hill-Cawthorne, Frances Bolt, Alison Holmes, Pantelis Georgiou, Jesus Rodriguez-Manzano

## Abstract

Real-time digital PCR (qdPCR) coupled with artificial intelligence has shown the potential of unlocking scientific breakthroughs, particularly in the field of molecular diagnostics for infectious diseases. One of the most promising applications is the use of machine learning (ML) methods to enable single fluorescent channel PCR multiplex by extracting target-specific kinetic and thermodynamic information contained in amplification curves. However, the robustness of such methods can be affected by the presence of undesired amplification events and nonideal reaction conditions. Therefore, here we proposed a novel framework to filter non-specific and low efficient reactions from qdPCR data using outlier detection algorithms purely based on sigmoidal trends of amplification curves. As a proof-of-concept, this framework is implemented to improve the classification performance of the recently reported ML-based Amplification Curve Analysis (ACA), using available data from a previous publication where the ACA method was used to screen carbapenemase-producing organisms in clinical isolates. Furthermore, we developed a novel strategy, named Adaptive Mapping Filter (AMF), to consider the variability of positive counts in digital PCR. Over 152,000 amplification events were analyzed. For the positive reactions, filtered and unfiltered amplification curves were evaluated by comparing against melting peak distribution, proving that abnormalities (filtered out data) are linked to shifted melting distribution or decreased PCR efficiency. The ACA was applied to compare classification accuracies before and after AMF, showing an improved sensitivity of 1.18% for inliers and 20% for outliers (p-value < 0.0001). This work explores the correlation between kinetics of amplification curves and thermodynamics of melting curves and it demonstrates that filtering out non-specific or low efficient reactions can significantly improve the classification accuracy for cutting edge multiplexing methodologies.

## INTRODUCTION

This manuscript demonstrates that undesired amplification reactions from real-time digital PCR (qdPCR) can be detected and filtered out by only evaluating the sigmoidal shape of an amplification curve. Here, we propose a novel methodology that can be used with multiplex PCR assays without the need of post-amplification analysis, increasing results accuracy and reliability.^1,2^

During the last decade, gold standard PCR technologies along with other nucleic acid amplification chemistries have resulted in key procedures for molecular diagnostic in both academic and clinical environments.^3–7^ However, limitations such as sample availability, trained personnel, and overall laboratory costs can represent obstacles to the scalability and adoption of PCR-based approaches.^8,9^ To overcome these barriers, multiplexing has been used to unlock the potential of conventional instruments, increasing the number of targets that can be detected in a single reaction.^10–12^ Since the adoption of multiplexing techniques, researchers and industries have successfully applied them to different areas such as molecular diagnostics, RNA signature polymorphism, and quantitative analysis.^13^ Moreover, in an effort to increase overall multiplex PCR capabilities, several studies have recently been published on the use of machine learning (ML) to identify the biological nature of an amplification event, improving throughput, clinical and analytical reliability, and sample classification accuracy.^14,15^ As described by Athamanolap et al. in 2014, ML methods were applied to High-Resolution Melt Curve to increase both the tolerance of melting temperature (*T_m_*) deviation among targets and reliability of classification for genetic variants (such as polymorphic genetic loci).^16^ In Jacky et al. 2021, ML techniques were used to enable high-level multiplexing using TaqMan probes by leveraging on single-feature classification (i.e. final fluorescence intensity or FFI) and PCR platforms with multiple fluorescent channels.^17^ On the other hand, Moniri et al. in 2020 proposed a new approach called Amplification Curve Analysis (ACA) for single channel multiplexing without explicitly extracting features. The ACA method comprises a supervised ML classifier to analyze kinetic information encoded in the entire amplification curve, by looking into sigmoidal shapes across different targets.^18^ Furthermore, using ACA along with Melting Curve Analysis (MCA), a new method called Amplification and Melting Curve Analysis (AMCA) was developed, enabling higher-level multiplexing in a single channel. The AMCA couples both ACA and MCA coefficients to improve classification accuracy. This has been demonstrated through the detection of nine mobilized colistin resistance genes and clinical isolates containing five common carbapenemase resistance genes.^19,20^

A barrier to wider adoption of the aforementioned techniques is that they may be limited by instrumentation specifications such as thermal profile performance, available optical channels/filters, and software setup. For example, MCA methodologies are particularly limited in combination with point-of-care devices and probe-based chemistries (such as TaqMan), as melting curve analysis is commonly not compatible. In these circumstances, the ACA method still stands as a valid option for multiplexing and therefore it has been the methodology of choice for the work proposed in this manuscript. However, across all these ML-based multiplexing strategies, the ACA approach can be negatively affected by the presence of abnormal amplification products, due to primer dimerization, amplification of undesired targets, miscalibration of the instrument, and intra-molecule secondary structures. These abnormal behaviors tend to alter the kinetic information of the sigmoidal curves, causing low efficiency or delaying the amplification reaction.^21–23^

As represented in **Figure 1**, when considering shapes of amplification curves from a multiplex assay, similarities among different targets can reduce the accuracy of the ACA classifier, as the presence of nonspecific or low efficient reactions results in blurred boundaries among clusters. To overcome this problem, we developed an intelligent algorithm to filter out outliers from multiplex amplification events. Furthermore, to validate the correctness of outlier removal, amplification curves (*inliers* and *outliers*) are compared with labeled melting curves (*correct* and *wrong*).

**Figure 1.**
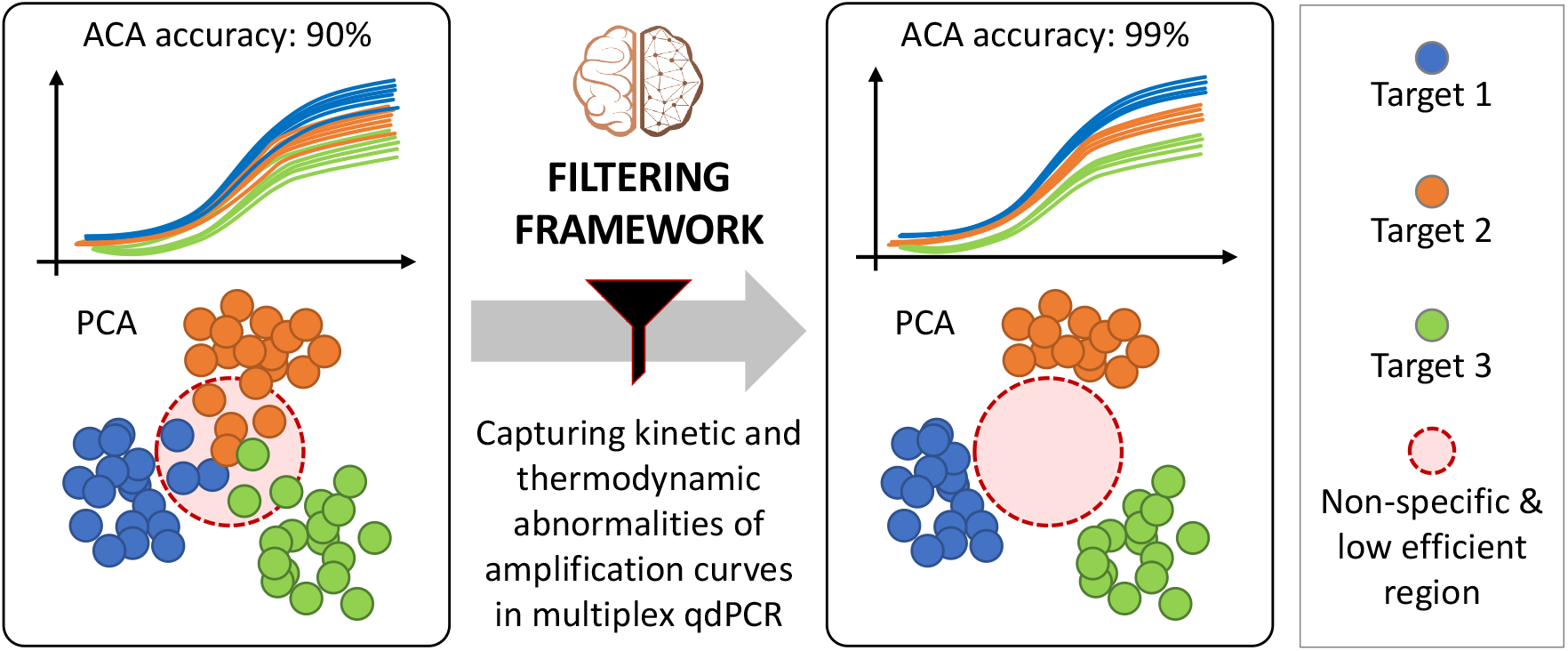
Concept figure. Left: raw amplification curves and their corresponding ACA clusters (represented by principal component analysis or PCA) include non-specific and low efficient reactions (confined in the red-circled region). The presence of outliers blurs the boundaries of the different clusters, negatively impacting ACA classification accuracy. By applying the proposed filtering framework, kinetic and thermodynamic abnormalities from amplification events can be captured. Right: Outliers are removed from the original data, resulting in more separated clusters and clearer boundaries. Therefore, ACA classification accuracy is improved.

In this work, we demonstrated that non-specific and low efficient PCR reactions affect the shape of the amplification curve and therefore, they can be filtered out considering only the sigmoidal trend. Furthermore, we developed an outlier removal algorithm called Adaptive Mapping Filter (AMF), which was used to improve the classification accuracy of the ACA method. These concepts were explored using data obtained from qdPCR experiments reported by Miglietta et al, 2021.^20^ As a case study, three of the “big 5” carbapenemase resistant genes (*bla_NDM_, bla_IMP_*, and *bla_OXA-48_*) were considered in this study.

Our vision is that by sharing this new approach we can significantly improve the quality of data from qdPCR instruments and enhance the sensitivity and accuracy of ML-based multiplexing methods relying only on amplification curves. Moreover, extending this framework to other amplification chemistries and real-time platforms will improve multiplexing capabilities of existing diagnostic workflows and platforms.

## EXPERIMENTAL SECTION

In this section, a new framework for outlier removal in qdPCR is proposed. As depicted in **Figure 2**, this framework took raw amplifica-tion curve data as input, and applied baseline and flat/late curve removal in the processing step. Then each processed curve was fitted by a sigmoid function and the fitted parameters, as well as a newly developed feature referred as *S_end_*, were used as input for a filtering algorithm which identified outliers automatically. Finally, the framework output the filtered amplification curves, marked as inliers.

**Figure 2.**
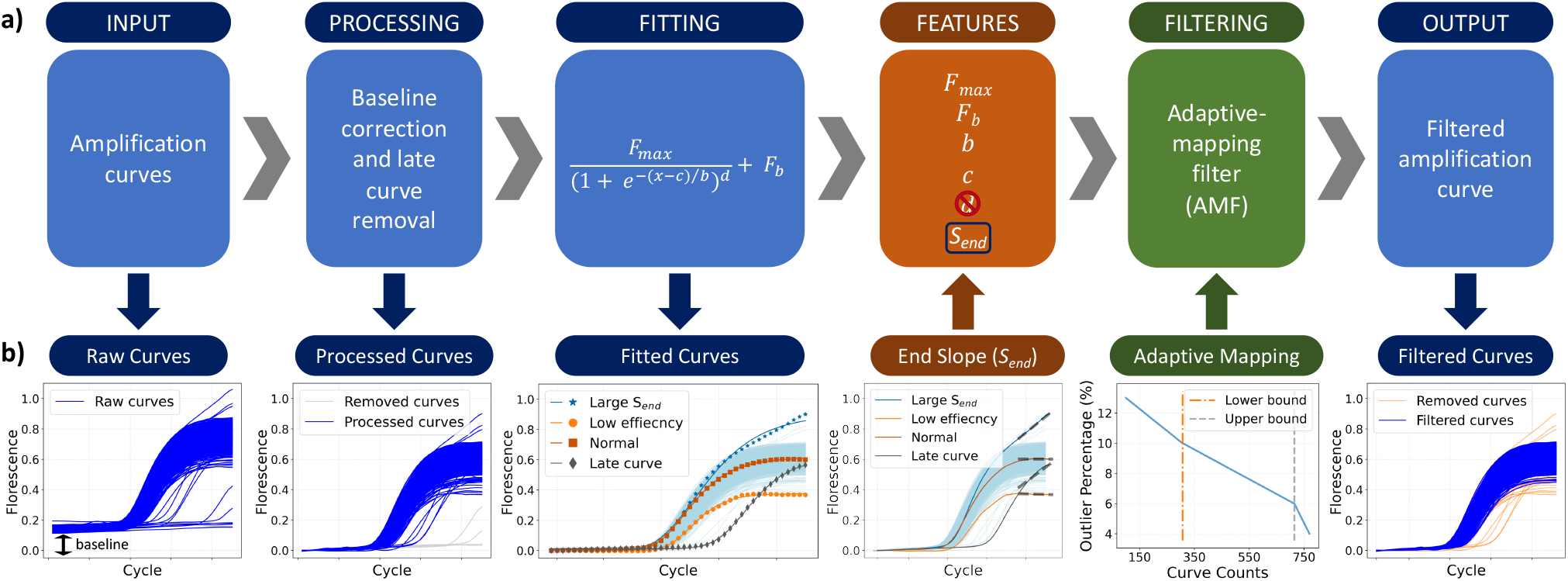
Proposed framework. a) Framework steps: raw data input, processing, curve fitting, feature extraction, Adaptive Mapping Filtering (AMF) and filtered curve output. b) Input or output of each step. From left to right, the input of the framework were raw amplification curves, some of which are flat or late curves. By applying the processing step, the baselines were removed, and flat/late curves were discarded. Following this, the processed curves were fitted using a five-parameter sigmoid function, after which each curve was condensed into five features. A new feature *S_end_* plus four of the parameters were used to form a set, which is the input of the filtering step. The *d* parameter was discarded from the feature set for filtering as it is unsuitable for the used algorithms. We further developed the AMF with a monotonic decreasing map between positive curve numbers within a panel and the outlier percentage. The outputs of the framework are the filtered curves (inliers).

### Data input

As a case study, data from Miglietta et al. 2021 was used in this work.^20^ Data from synthetic DNA (gBlocks™ gene fragments, IDT) containing *bla_NDM_* (N=18,480), *bla_MP_* (N=17,710), and *bla*_OXA-48_ (N=17,710) gene sequences were used as the training dataset, and 198 clinical isolates labeled with these three targets were used as the testing samples. Each sample contained 770 raw curves for a total of 152,460 curves across all the samples, within which 116,222 were positive after the processing step. It is expected that data from clinical isolates are much noisier and thus contain more outliers than those from gBlocks.

### Data processing

The first step of the framework is processing the raw curves using a baseline correction and a flat/late curves removal to exclude the negative curves of the unprocessed data from the qdPCR output. The baseline of real-time PCR reaction during the initial cycles presents little change in fluorescent signal. The low-level signal of the baseline equates with the background or noise of the reaction. Therefore, we processed the baseline of each raw curve by averaging the fluorescent value of the first five cycles and subtracting it from the time series. Following this, flat/late curves were removed by applying an upper and lower fluorescence threshold at the 40^th^ cycle, as suggested by the manufacturer.^24,25^

### Fitting and feature extraction

Following the processing step, a curve fitting step was introduced to represent the processed amplification curves with sigmoid parameters, which were later down-selected and used as input features for outlier removal and classification algorithms. A 5-parameter sigmoid model,^23^ which is shown below, was used to fit the amplification curves:

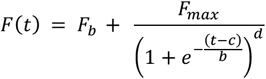

where *t* is the PCR cycle number, *F*(*t*) is the fluorescence at the *t*^th^ cycle, *F_b_* is the background fluorescence, *F_max_* is the maximum fluorescence, *b* relates to the slope of the curve, *c* is the fractional cycle of the inflection point, and *d* is the asymmetric parameter. To solve the nonlinear least-square optimization problem for the curve fitting, the Trust Region Reflective (TRF) algorithm with specific bounds was used.^26^ Here, we set [10, 0.3, 10, 50, 100] and [0,-0.1,-10,-50,-10] as the upper and lower bounds for the search of the 5-parameter set *p* = [*F_max_, F_b_, h, c, d*]. The initial parameter set *p*_0_ was optimized through pivot fitting on 5% of the training data. After fitting, each amplification curve was given as five parameters, which are condensed representations of curve information. All parameters except for *d* were considered as input features for outlier removal algorithms because parameter values of outliers may have significant differences from those of normal curves. The *d* parameter shows a bimodal distribution with 2 distant peaks, which is unsuitable for the outlier removal step because many of the outlier algorithms require a unimodal distribution of features. Therefore, the *d* parameter was discarded from the feature set for filtering.

In addition, we further introduced a new feature called the end slope (*S_end_*) with the aim to provide further information about the amplification curve shape. This was calculated by taking the average of the first derivatives at the last five cycles of the amplification curve:

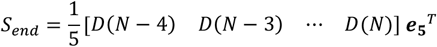

where:

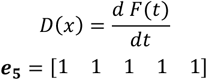

and *N* is the total cycle number.

Using the *S_end_* feature, the information in the tail of amplification curves was extracted, which contributes to distinguishing inliers and outliers. For example, as illustrated in the “Fitting Curves” step of **Figure 2b**, curves that don’t reach the plateau may have larger end slopes. These curves cannot be precisely represented by the fitted parameters since the fitting equation is not capable to capture this non-plateaued trend. Therefore, *S_end_* would benefit the result of outlier removal by providing additional information to the feature set. Including *S_end_* and discarding *d*, the final feature set for outlier removal algorithms is ***x_f_*** = [*F_max_, F_b_, b, c, S_end_*].

### Outlier removal algorithms

In this research, seven outlier removal algorithms were considered, which can be split into the following categories according to their principal ideas of filtering: proximity-based, linear, outlier ensembles, and angle-based algorithms. (i) Proximity-based outlier detection algorithms rely on using a distance metric (e.g. Euclidean or Manhattan) to identify outliers. We applied two proximity-based algorithms which are Local Outlier Factor (LOF) and Density-based Spatial Clustering of Applications with Noise (DBSCAN).^27,28^ The LOF algorithm considers the *k*-nearest neighbors (KNN) to every point in the dataset and computes a local outlier factor for each of them. DBSCAN classifies the points into the core, border, and noise of clusters based on the number of points (min points) within the radius (epsilon) of the considered point. (ii) The linear outlier detection methods used were One-Class Support Vector Machine (OC-SVM) and Elliptical Envelope.^29,30^ OC-SVM applies the concept of finding a hyperplane that separates the inlier points from the origin, such that the hyperplane is closest to the inlier points as possible. The Elliptical Envelope aims to fit the smallest ellipse possible to the core cluster of data points, with any point outside being considered outliers. (iii) Outlier ensemble-based detection methods considered were Isolation Forest and feature bagging.^31,32^ Isolation Forest uses random forests to recursively randomly partition data, after which datapoints with fewer partitions to isolate are marked as outliers. Feature bagging considers multiple outlier algorithms and randomly selects a group of features. From those features, the resulting outlier scores from each algorithm are merged to find the strongest outliers. (iv) Angle-based Outlier Detection considers the angles made by a point with all other pairs of points in the dataset.^33^ For each point, the variance is calculated from all the angles obtained, where for a potential outlier the variance is small, since the point is distant from the main cluster of data.

### Adaptive mapping filter (AMF)

Most of the outlier detectors explained in the previous section require a hyperparameter called “contamination ratio” or “outlier percentage”, which represents the percentage of outliers to be removed from the original data. To adaptively set up this hyperparameter, we developed a mapping strategy that maps the number of positive reactions per panel in the qdPCR chip (processed curves) to the contamination ratio used in the outlier removal algorithm.

In digital PCR, as the number of positive curves increases, the probability of having more than one molecule in a single well increases, resulting in a shift of reaction state from digital to bulk. Moreover, as the reaction goes toward the bulk region, a higher number of positive curves will be present in a panel, which can result in a lower probability of observing a non-specific or low efficient reaction (outlier) in a well.^18,34^ Let us suppose that for each well the probability of observing an outlier is *p*(*M_i_*), where *M_t_* is the number of processed curves for the *i* th sample. Since *p*(*M_i_*) are independent and identical distributed (*i.i.d*.) for all the wells, the total number of outliers *X_i_* observed in the *i*^th^ sample follows the distribution of *X_i_~B*(*M_i_, p*(*M_i_*)). Therefore, the expected percentage of outliers in the *i*^th^ sample should be:

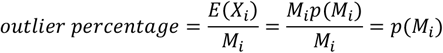

which means that the expected outlier percentage is a monotonic decreasing function to the number of positive curves. In this research, we applied a piecewise linear function with empirical turning points, as illustrated in the filtering step of **Figure 2b**.

Coupling the adaptive mapping with an outlier removal algorithm, we developed a novel method called Adaptive Mapping Filter (AMF), which takes as input the feature set and output the inliers.

### Melting Labeling

An algorithm was developed to automatically label the melting curves as specific (which we called “correct”) or nonspecific (referred as “wrong”) ones. By using this methodology, the percentage of wrong melting curves within all the curves of a sample (Wrong Melting Percentage or WMP) was calculated, and this WMP further served as a metric for performance evaluation.

To apply melting labeling, the reference melting peak for each target needs to be determined. For a target *tg* ∈ |*bla*_NDM_. *h/a*_IMP_ *h/a*_OXA-48_], a reference melting peak temperature 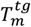 was given by calculating the median value of all the melting peak temperatures of the gBlock curves with target *tg*. After that, the steps below were followed to label every single melting curve of the clinical dataset:

1. Find the global maximum melting peak’s temperature 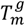 of the current melting curve.
2. If 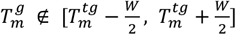, where *W* is the tolerance width of the 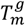 distribution, the current curve is labeled directly as a wrong melting curve.
3. Otherwise, find the local maximum melting peaks’ temperatures on the left and right sides of 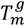 on the current curve, mark them as 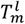 and 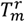 respectively. Note that either 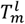 or 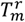 may not exist. If neither exists, the current curve will be labeled as a correct melting curve.
4. If at least one of 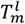 and 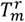 exists, a set of this (these) local melting peak(s) will be constructed. For each element 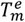 in this set, check whether

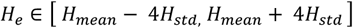

where *H_e_* is the height of the current melting curve at temperature 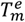 and *H_std_* are the mean and standard deviation of 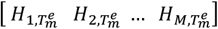, in which 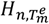 means the height of the n^th^ melting curve of the sample at temperature 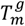, and *M* is the total curve number in the sample. If at least one of the above tests fails, the current curve will be labeled as a wrong melting curve. Otherwise, it will be marked as a correct one.

With the above steps, it is ensured that both curves with large deviations of 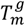 from reference melting peaks and curves with large nonspecific local melting peaks can be labeled as wrong. In this way, all the curves had been marked as either “correct” or “wrong”, and further used to calculate the Wrong Melting Percentage (WMP):

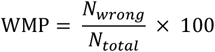

where *N_wrong_* is the number of wrong melting curves within the sample, and *N_total_* is the total number of curves in the sample.

It is worth mentioning that the proposed algorithm of automatic melting labeling is not a part of the filtering framework. The labeling was used to calculate the WMP which functioned as a metric for filtering evaluation, where a lower WMP indicates better filtering performance.

### Data visualization

Visualization is a vital step for understanding the distribution of a given dataset. In this article, Principal Component Analysis (PCA) with 2 components was used to visualize the feature sets of the curves before and after applying the outlier removal algorithm into scatter plots. Visual inspection was performed to illustrate how separated the clusters of different targets were. Following this, several metrics for measuring density and degree of separation among those clusters were used to quantitively evaluate how well they were divided.

Specifically, after the PCA of the feature set ***x_f_*** = [*F_max_, F_b_, b, c, S_end_*] from the amplification curves of each target, the Silhouette Coefficient for each feature set was calculated.^35^ The mean value of these coefficients, known as the mean Silhouette Score, was then used to indicate how well the curves of the same targets are clustered. A Higher Silhouette Score implies denser and better-separated clusters observed. Two additional metrics, the Calinski-Harabasz score and the Davies-Bouldin score, were also implemented for clustering evaluation, where a higher Calinski-Harabasz score or a lower Davies-Bouldin score relates to larger inter-cluster distances among targets.^36,37^

### Classification of amplification curves – data-driven multiplexing

The ACA method uses kinetic information encoded in the amplification curve to classify different nucleic acid molecules from a PCR test. To illustrate the influence of the AMF on the ACA, a random forest classifier with 100 trees was applied to the feature set ***x_c_*** = [*F_b_, F_max_, b, c, d*], which differs from the ***x_f_*** used for outlier removal algorithms. Here, parameter *d* was reintroduced because more curve-related information is needed, provided that the proposed classifier is relatively less sensitive to the feature distributions. *S_end_* was discarded for classification because, after outlier removal, abnormal curves with large end slopes were not present in the data set. For the remaining curves, *S_end_* were extremely close to zero, thus it was not necessary for *S_end_* to be included again. All the other features were normalized with the mean and the variance of the training data before being input into the classifier.

For the ACA, processed gBlock data were used as the training dataset, as mentioned in the “Data Input” section. For the testing set, we utilized both the inliers and the outliers marked by the aforementioned AMF algorithm and tested them respectively. As a comparison, two randomly down-selected datasets with the same numbers of curves as the inliers and the outliers were also constructed and tested.

### Statistical Analysis

Two-sided Wilcoxon signed-rank tests were used to determine the statistical significance of the changes of WMP and melting peak distributions (distributions of melting peak temperature, *T_m_*, and height, *H_m_*) before and after outlier removal. Two-sided Mann-Whitney U rank tests were used to compare the distributions of *C_t_*, FFI, and maximum slopes between inlier and outlier amplification curves. Those three metrics were chosen for their relationship with the amplification curve efficiency. Many studies suggest that sigmoidal modeling of the entire amplification curve can be used to define the rate of the PCR efficiency. Therefore, low efficient PCR reactions are related to low fluorescent values and low maximum slope.^38,39^

Moreover, the significance of the comparison between inliers and outliers in clustering Silhouette coefficients was determined by a twosided Wilcoxon signed-rank test. This test was also used in the evaluation of the classification performance. A p-value of 0.001 with Bonferroni correction was used as the threshold for statistical significance.

## RESULT AND DISCUSSION

In this study, a new framework is presented to detect outliers from amplification reaction in qdPCR. The outlier identification relies on the AMF, which is comprised of an outlier detection algorithm and a mapping strategy to adapt the contamination ratio hyperparameter to the positive amplification reaction counts (or positive wells) of the qdPCR chip.

### Evaluation of outlier detection algorithms

As shown in **Figure 3a**, we evaluated the detection performance of seven outlier removal algorithms on filtering amplification curves against outlier percentages by using three metrics: (i) Wrong Melting Percentage (WMP), (ii) Melting Curve *T_m_* variance, (iii) Melting Curve *H_m_* variance. The changing values of metrics for different algorithms with fixed outlier percentages from 0.1% to 40% are shown in **Figure 3a**. After the filtering is applied, the WMP shows a significant reduction from 1.1% (from the unfiltered dataset) to a maximum of 0.9% after filtering across all the algorithms. The graph depicts that outlier percentage and WMP are inversely proportional, but the trend can vary among methods. Proximity-based outlier detectors perform worse overall compared to the rest so they are unable to achieve a dramatic decrease in WMP, even with very large contamination ratios. On the other hand, ensemble-based detectors such as Feature Bagging and Isolation Forest have better performance with the lowest WMP among all the outlier percentages. As shown in the center and right end graphs, the variances of *T_m_* and *H_m_* have a decreasing trend that can be observed as the outlier percentage increases, indicating that both of their distributions are narrowed down. In the *T_m_* variance plot, it is noticed that DBSCAN achieves better performance at lower outlier percentages, but this trend reaches a plateau as the outlier percentage further increases. Once again, ensemble-based methods have similar behavior for the *T_m_* variance as for the WMP. For instance, Isolation Forest outperforms all other detectors after the outlier percentage reaches 12%. Moreover, Isolation Forest and elliptic envelop show the best performance for *H_m_* variance up to 26% contamination ratio.

**Figure 3.**
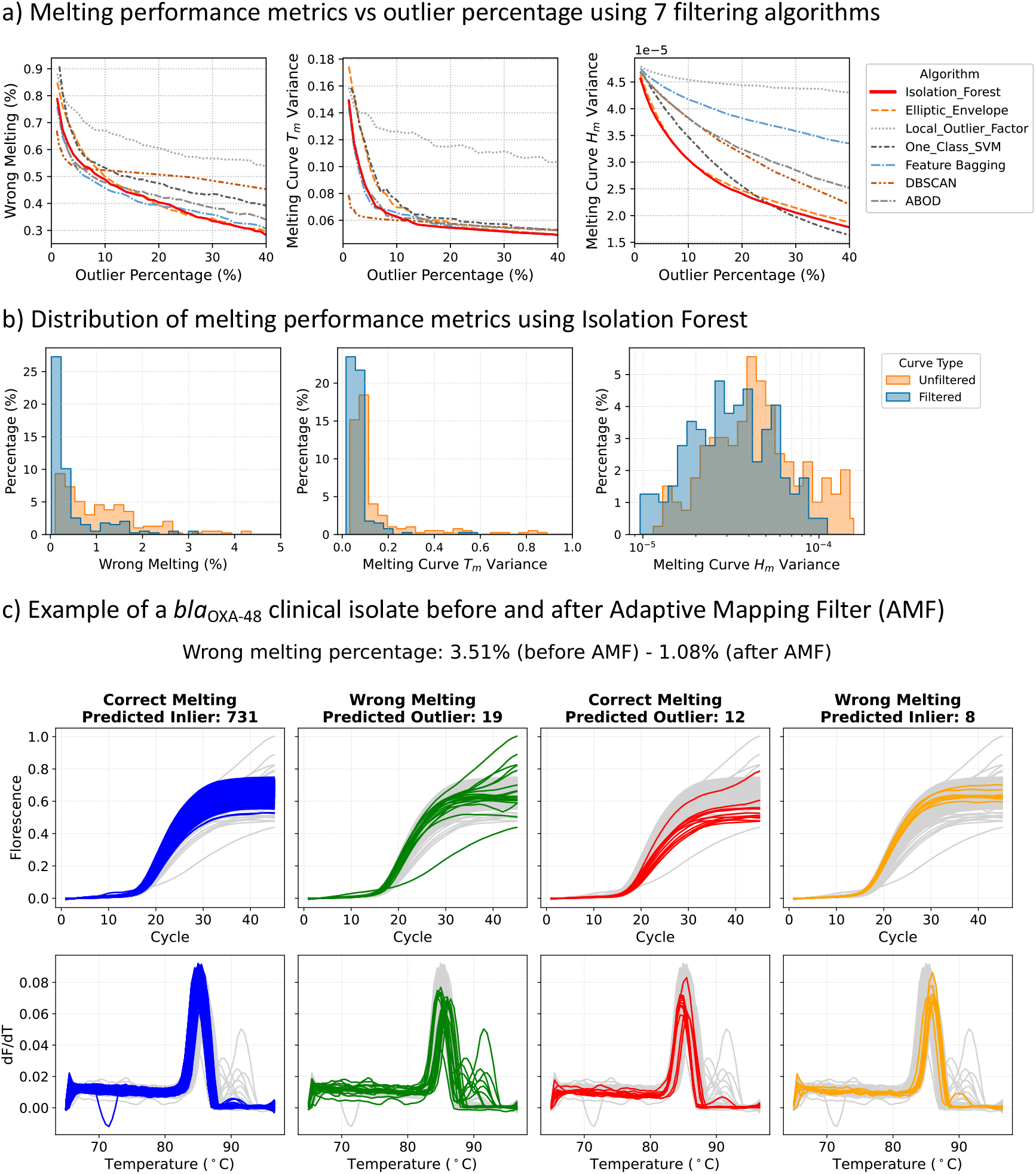
Melting curve analysis on filtered results. a) Melting performance shown with Wrong Melting Percentage, *T_m_* and *H_m_* variances versus fixed outlier percentage. As the outlier percentage increases, all the metrics show decreasing trends which tend to plateau after a certain percentage. As illustrated by the firm red line, Isolation Forest performs the best overall for the three metrics. b) The distribution of melting performance metrics shows that, after filtering, the WMP becomes significantly smaller, and *T_m_* and *H_m_* have a narrower distribution. c) An example of *bla*_OXA-48_ clinical isolate. Each column shows the amplification curve and corresponding melting curve of the correct melting and predicated inliers (N=731), wrong melting and predicted outliers (N=19), correct melting and predicted outliers (N=12), wrong melting and predicted inliers (N=8).

In this analysis, WMP was used to show the change of wrong melting proportion after applying outlier detection algorithms, indicating the direct effect of the filtering on removing wrong melting curves. Moreover, a smaller *T_m_* variance indicates a narrower *T_m_* distribution, which in combination with the WMP methods shows that curves with large deviations from the reference 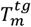 are removed by the filtering algorithm. In molecular biology, those curves may be generated after non-specific events such as undesired target interaction or primer dimerization.^40^ In addition, melting curves presenting low-*df/dt* (or *H_m_*) are associated with low efficient amplification reactions. Therefore, narrowed distribution of *H_m_* indicates that low efficient curves, which are present at the tail of the distributions, are removed.^41^ All the algorithms provide better performance compared to the original benchmark calculated on the unfiltered data. However, it is noticed that Isolation Forest is always among one of the best methods for all the metrics and does not show any defects, which is common for other algorithms. In the following sections, we use Isolation Forest to further demonstrate the proposed framework.

### Filtering performance analysis of the AMF

In the following step, AMF was applied to the unfiltered data, and the distributions of inner-sample WMP, *T_m_* and *H_m_* variances are illustrated in **Figure 3b**. Across these three metrics, significant shifts of distributions to smaller values are shown after filtering, supported by all the p-values < 0.0001. This indicates that the proposed AMF can significantly remove both non-specific and low efficiency curves only by looking at amplification curves. This proves our hypothesis that amplification curves contain not only kinetic but also thermodynamic information as numbers of filtered outliers correspond to wrong melting curves.

An example of the AMF visual performance on a clinical isolate containing the CPE gene *bla*_OXA-48_ is illustrated in **Figure 3c**. Columns represent both amplification and melting curves of: (i) correct melting and predicated inliers (N=731, 94.9%), (ii) wrong melting and predicted outliers (N=19, 2.5%), (iii) correct melting and predicted outliers (N=12, 1.6%), (iv) wrong melting and predicted inliers (N=8, 1%). The first column shows the correctly identified inliers representing specific products of PCR tests. In the second column, non-specific reactions are correctly identified and labeled as outliers, which emphasizes the effectiveness of the filtering. We noticed that a small number of specific curves were predicted as outliers, as shown in the third column of **Figure 3c**. This phenomenon does not deny the efficacy of the filter, as these “incorrectly” removed curves have: (i) significantly larger *C_t_* values, (ii) significantly smaller FFI, (iii) and smaller values of maximum slope compared to the inliers. Across the entire clinical isolate dataset (N=116,222), compared to melting curve analysis, 115,535 were correctly predicted inliers and 791 were correctly predicted outliers. Furthermore, 5,861 were wrongly classified as outliers whereas 687 were wrongly classified as inliers. Further statistical analyses on the entire dataset also endorse these significant differences between inliers and outliers for *C_t_*, FFI and maximum slope values, as illustrated in **Table 1**. This indicates that AMF removes certain curves because they are of low amplification efficiencies even though they have “correct” melting peaks. A few curves labeled as “wrong” melting may be predicted as inliers, as shown in the fourth column of **Figure 3c**. This can be explained by the relatively low temperature resolution of the equipment which results in mislabeled wrong melting curves due to the large quantization noise of 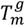 during temperature measurement. In fact, by visually inspecting the last column of **Figure 3c**, it can be seen that amplification curves are of very similar shapes to correctly predicted inliers (shown in the first column of **Figure 3c**). The WMP of the illustrated sample has dropped from 3.51% to 1.08%.

**Table 1.** Comparison of *C_t_*, FFI and maximum slope between predicted inliers and outliers with correct melting peaks.

| Target | $C_t$ (mean ± std) | | FFI (mean ± std) | | Max Slope (mean ± std) | |
| --- | --- | --- | --- | --- | --- | --- |
|  | Inliers | Outliers | Inliers | Outliers | Inliers | Outliers |
| *bla<sub>NDM</sub>* | 21.45 ± 3.28 | 26.40 ± 6.11 | 0.67 ± 0.06 | 0.60 ± 0.12 | 0.07 ± 0.01 | 0.06 ± 0.01 |
| *bla<sub>IMP</sub>* | 30.33 ± 2.23 | 31.25 ± 3.43 | 0.44 ± 0.06 | 0.41 ± 0.07 | 0.0276 ± 0.003 | 0.0271 ± 0.01 |
| *bla<sub>OXA-48</sub>* | 18.82 ± 3.08 | 21.03 ± 4.34 | 0.65 ± 0.08 | 0.51 ± 0.16 | 0.05 ± 0.01 | 0.04 ± 0.02 |

For all the targets, inliers have significantly smaller $C_t$ and larger FFI and max slope, with all $p$ -values $< 0.0001$ .

### Feature sets visualization

To visualize the effect of the AMF, PCA-based feature visualization before and after filtering is depicted in **Figure 4**. On the left of the figure, the unfiltered data shows larger overlapping within clusters of different targets and a higher number of outliers compared to the filtered data. The segmented squares are used to emphasize the differences in cluster overlapping before and after the AMF, where clearer boundaries between *bla*_IMP_ and both *bla*_OXA-48_ and *bla*_OXA-48_ can be seen. These differences highlight that: (i) outliers can be effectively removed by the AMF, and (ii) removing outliers enhance the separation and reduce the overlap among different target clusters, which will ease the classification of the ACA method.

**Figure 4.**
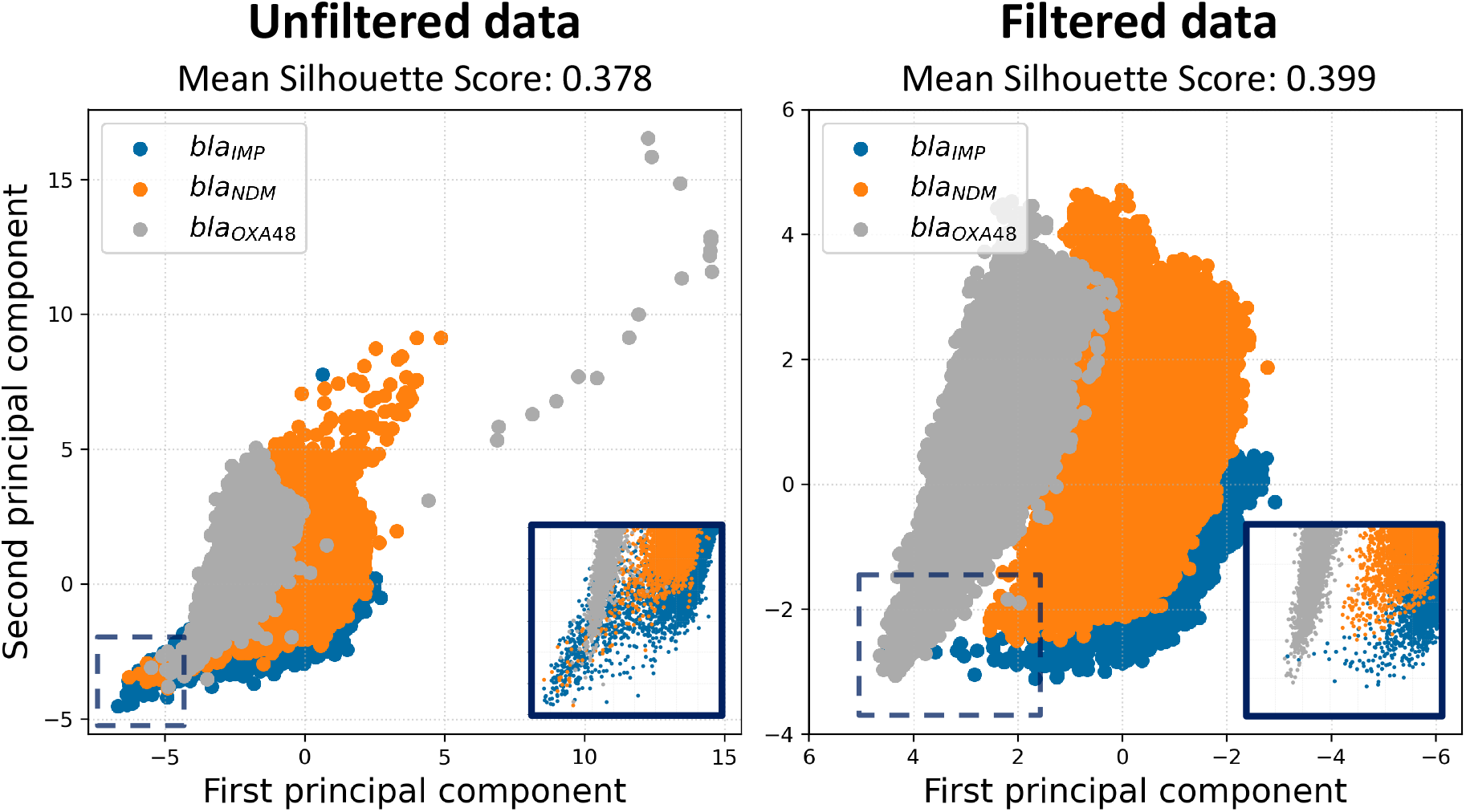
Data visualized using 2-D Principle Component Analysis before and after filtering. The filtered data plot shows that most outliers have been removed from the original unfiltered data, and the clusters are more separated with clearer boundaries and fewer overlaps. The segmented squares on the bottom side of both figures show the areas where cluster overlapping is more evident, thus they are zoomed. The mean Silhouette Score rises from 0.378 to 0.399 after filtering.

To numerically evaluate the degree of separation across target clusters, the mean Silhouette score of all the datapoints was calculated before and after filtering, showing an increment from 0.378 to 0.399 (p-value < 0.0001). In addition, the Calinski-Harabasz score increased from 101,002.729 to 130,134.802, and the Davies-Bouldin score dropped from 0.886 to 0.839. All those results indicate that AMF makes target clusters denser and better separated.

### ACA classification

After demonstrating that removing outliers improves the overall distance among clusters, we further explored its impact on the ACA classification for both inlier and outliers against randomly down-selected datasets with the same numbers of curves. In **Figure 5a**, the confusion matrix shows that the sensitivity for the inliers is 88.96%, which is an increase of 1.13% compared to the randomly down-selected ones (**Figure 5b**). For all the targets, a significant sensitivity improvement can be observed of 1.06%, 0.95% and 1.39% for *bla*_IMP_, *bla*_NDM_, and *bla*_OXA-48_, respectively. Moreover, the overall classification accuracy was 84.94% for inliers and 83.76% for randomly down-selected curves, showing a 1.18% improvement (*p*-value < 0.0001), which is in line with the overall WMP before filtering (WMP=1.1%). Applying the filter will help increase the overall performance and specificity of the dataset. This supports our hypothesis that melting information or thermodynamics are contained in the amplification curve.

**Figure 5.**
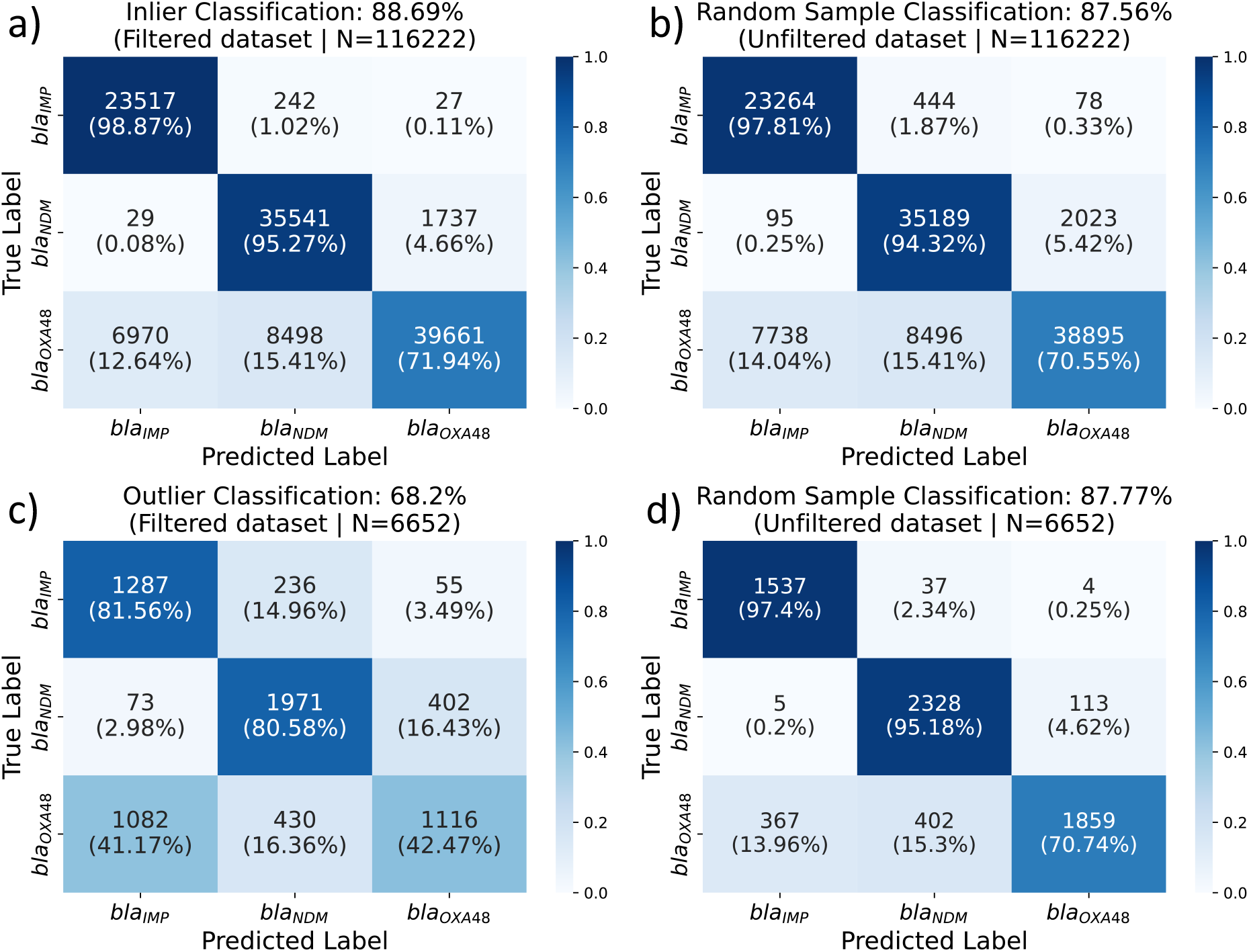
Confusion matrices for inlier and outlier classification. The four confusion matrices are shown for: a) inliers, b) randomly down-selected data with the same curve numbers as inliers, c) outliers, and d) randomly down-selected data with the same curve numbers as outliers. The title of each matrix reports the sensitivity of the model. Moreover, each square of the matrix has the number of predicted curves for the corresponding true label and the respective sensitivity of the square.

To show that the removed outliers are less informative for targets recognition and harmful for the overall classification, in **Figure 5c-d** we show the confusion matrices of the classification using both removed outliers and a randomly down-selected dataset with the same size. As expected, the performance for outliers is significantly worse than the randomly down-selected ones, with only 68.2% and 54.78% sensitivity and accuracy respectively for outliers (*p*-values < 0.0001). This dramatic sensitivity decrement of 19.57% strongly suggests that outliers have less useful information for the classification of the selected targets.

Furthermore, the statistical analysis on the two randomly down-selected datasets shows no significant differences of in-sample accuracy with *p*-value = 0.448, which is in line with the central limit theorem as they originate from the same distribution. This is a further proof that the efficacy of the proposed framework is not related to the size of the data.

## CONCLUSION

In this paper, we presented a novel framework to adaptively remove abnormal curves from PCR amplification reactions. The method takes the raw input from a qdPCR run and processes it in three steps: background subtraction, late curve removal, and sigmoidal fitting. Moreover, a new feature called end slope (*S_end_*) is developed in this study which, along with sigmoidal parameters, is used in the Adaptive Mapping Filter (AMF). The AMF is capable of removing non-specific and low efficient amplification curves, which are labeled as outliers. Melting curves of the outliers, previously removed, were compared with melting curves of inliers using both Wrong Melting Percentage (WMP) and melting peak distributions. Results show that non-specific and low efficient curves can be removed from amplification reaction by purely considering the sigmoidal trend. Further validation of the framework performance was conducted by assessing the classification accuracy and sensitivity of the ACA classifier on both inliers and outliers. This reinforces our hypothesis that removing abnormalities of amplification reaction in real-time PCR instruments would benefit data-driven multiplexing by removing undesired information.

In this research we used data from qdPCR published in our previous work to demonstrate the effectiveness of the proposed framework, but its generality has not been tested in other settings. Future work will focus on evaluating this methodology on real-time data originating from various qPCR instruments, from different chemistries (such as isothermal amplification), and from point-of-care devices. Moreover, upcoming work will focus on introducing advanced machine learning techniques to enhance the classification efficacy of the ACA classifier, and then on making this approach more reliable for use in clinical diagnostics.

In conclusion, this study reveals the interconnection between the kinetics of the amplification curve and the thermodynamics of the melting curves. For the first time, a framework is introduced which is capable of removing abnormalities in kinetic and thermodynamic information by purely screening amplification curves.

## Author Contributions

All authors have given approval to the final version of the manuscript.

## ACKNOWLEDGEMENT

This work was supported by the Imperial COVID-19 Research Fund (WDAI.G28059); the Department of Health and Social Care-funded Centre for Antimicrobial Optimisation (CAMO) at Imperial College London; the Imperial College President’s PhD Scholarships 2021 (KX). Authors KHC, FB, AH, PG and JRM are affiliated with the NIHR Health Protection Research Unit (HPRU) in Healthcare Associated Infections and Antimicrobial Resistance at Imperial Collzege London in partnership with the UK Health Security Agency (previously PHE) in collaboration with, Imperial Healthcare Partners, the University of Cambridge and the University of Warwick. The views expressed in this publication are those of the authors and not necessarily those of the NHS, the National Institute for Health Research, the Department of Health and Social Care, or the UK Health Security Agency. AH is a National Institute for Health and Care Research (NIHR) Senior Investigator.

## Insert Table of Contents artwork here

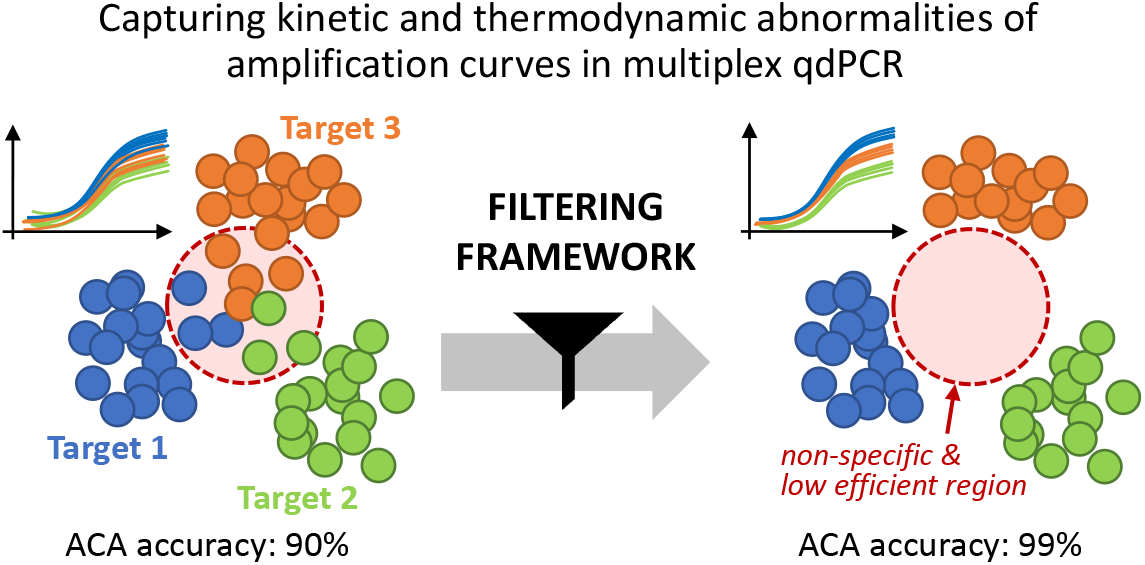

